# Bacteriocin Distribution Patterns in *Enterococcus faecium* and *Enterococcus lactis:* Bioinformatic Analysis Using a Tailored Genomics Framework

**DOI:** 10.1101/2023.11.13.566347

**Authors:** Ana P. Tedim, Ana C. Almeida-Santos, Val F. Lanza, Carla Novais, Teresa M. Coque, Ana R. Freitas, Luísa Peixe

## Abstract

Multidrug-resistant *Enterococcus faecium* represent a major concern due to their ability to thrive in diverse environments and cause life-threatening infections. While antimicrobial resistance and virulence mechanisms have been extensively studied, the contribution of bacteriocins to *E. faecium*’s adaptability remains poorly explored. *E. faecium*, within the Bacillota phylum, is a prominent bacteriocin producer. Here, we developed a tailored database of 76 Bacillota bacteriocins (217 sequences, including 40 novel bacteriocins) and applied it to uncover bacteriocin distribution patterns in 997 quality-filtered *E. faecium* and *Enterococcus lactis* (former *E. faecium* clade B) genomes. Curated using computational pipelines and literature mining, our database demonstrates superior precision versus leading public tools in identifying diverse bacteriocins. Distinct bacteriocin profiles emerged between *E. faecium* and *E. lactis*, highlighting species-specific adaptations. *E. faecium* strains from hospitalized patients were significantly enriched for bacteriocins as *ent*A, *bac*43, *bac*AS5 and *bac*AS11. These bacteriocins strongly associated with antibiotic resistance, particularly vancomycin and ampicillin, and Inc18 *rep*2_pRE25-derivative plasmids, classically associated with vancomycin resistance transposons. Our integrated genomic and epidemiological analysis elucidates meaningful connections between bacteriocin determinants, antimicrobial resistance, mobile genetic elements, and ecological origins in *E. faecium*. This work significantly expands the knowledge on the understudied bacteriocin diversity in opportunistic enterococci, revealing their contribution to environmental adaptation. Further characterization of strain-level bacteriocin landscapes could inform strategies to combat high-risk clones. Overall, these insights provide a framework for unravelling bacteriocins’ therapeutic and biotechnological potential.

## INTRODUCTION

Enterococci are ubiquitous in nature, as they constitute a natural component of the gastrointestinal microbiota in numerous mammals, including humans [1]. Nonetheless, they have also been linked to life-threatening infections, especially multidrug-resistant (MDR) *Enterococcus faecium* [1, 2]. *E. faecium* populations have evolved multiple adaptive features, enabling them to thrive in diverse environments, making them proficient colonizers in various niches and hosts [2]. While extensive research has explored antimicrobial resistance, virulence factors, and metabolic pathways in recent years, the role of bacteriocins in their adaptation to various niches, particularly the hospital, remains scarcely explored [3]. Within the Bacillota phylum, *E. faecium* stands out as one of the greatest producers of bacteriocins [2]. Bacteriocins, are ribosomally synthesized antimicrobial peptides that confer a competitive advantage in different environments, such as the gut microbial network, by selectively inhibiting the growth of particular pathogens [4]. Bacteriocins are structurally and genetically diverse, located on disparate gene clusters often located on plasmids alongside genes encoding antimicrobial resistance and virulence [2, 4].

While bacteriocin classification remains somewhat controversial, two well-defined major classes are lantibiotics (class I) and unmodified non-lantibiotics (class II) [2, 3, 5]. Bacteriolysins are usually considered separately [5]. The majority of known bacteriocins identified in enterococci belong to class II, which are small (<10kDa), heat-resistant peptides that do not undergo extensive post-translational modification as lantibiotics [2, 3]. They can be further divided into class IIa (e.g., enterocins A and P, and bacteriocin 43), class IIb (e.g., enterocin C, 1071 and X), class IIc (e.g., bacteriocin AS-48 and enterocin 4) and class IId (e.g., Enterocin Q and L50) [2, 3]. Some studies also consider a 3^rd^ class of bacteriocins, Class III, that includes bacteriolysins. These bacteriolysins cleave the peptidoglycan cross-links of the target cell wall causing cell death. For an in-depth understanding of enterococci*’* bacteriocins please refer to our recent review [3].

Currently, there are five databases specialized in bacteriocins: BACTIBASE [6], LABiocin [7], BAGEL [8], antiSMASH [9] and BUR (https://drissifatima.wixsite.com/bacteriocins). Among these databases, only BAGEL4 (http://bagel4.molgenrug.nl/) and antiSMASH (https://antismash.secondary metabolites.org/) offer direct bacteriocins searches in bacterial genomes. However, there is limited information available regarding the number of bacteriocins sequences and the bacterial species included in these databases. In light of these limitations and the undeniable relevance of bacteriocins in enterococci, we have developed a dedicated Bacillota bacteriocin database specifically designed for enterococci and used it to identify the presence of bacteriocins in 997 genomes of *E. faecium* and *Enterococcus lactis* (former *E. faecium* clade B) [10]. We also analysed the association between bacteriocins and antibiotic resistance genes, plasmid backbone genes and clones in order to elucidate the impact of bacteriocins in the adaptation of *E. faecium* and *E. lactis* to different environments. This new database will help to explore the presence and diversity of bacteriocins in enterococci linked to different Public Health contexts, offering valuable information about their ecological significance of potential therapeutic and biotechnological value.

## METHODS

### Bacterial genome collection

A total of 1,676 genomes classified as *E. faecium* were extracted from the GenBank/PATRIC database until January 16^th^, 2020, which included all available genomes up to 2019. Core genome Multilocus Sequence Typing (cgMLST) and Multilocus Sequence Typing (MLST) analysis was performed on all the 1,676 isolates, including *E. faecium* and potentially *E. lactis*, using Ridom SeqSphere+ v5.1.0 (Ridom GmbH, Münster, Germany), using the default settings of the program.

Strains with the same cgMLST cluster types (CTs) were considered as clones if they had originated from the same country/city, same isolation source, and isolated in the same year. Genomes lacking information regarding geographical location and/or date of isolation were considered similar clones if they belonged to the same study. After applying these criteria, our database ended with a total of 997 unique genomes (Figure 1.A).

**Figure 1.**
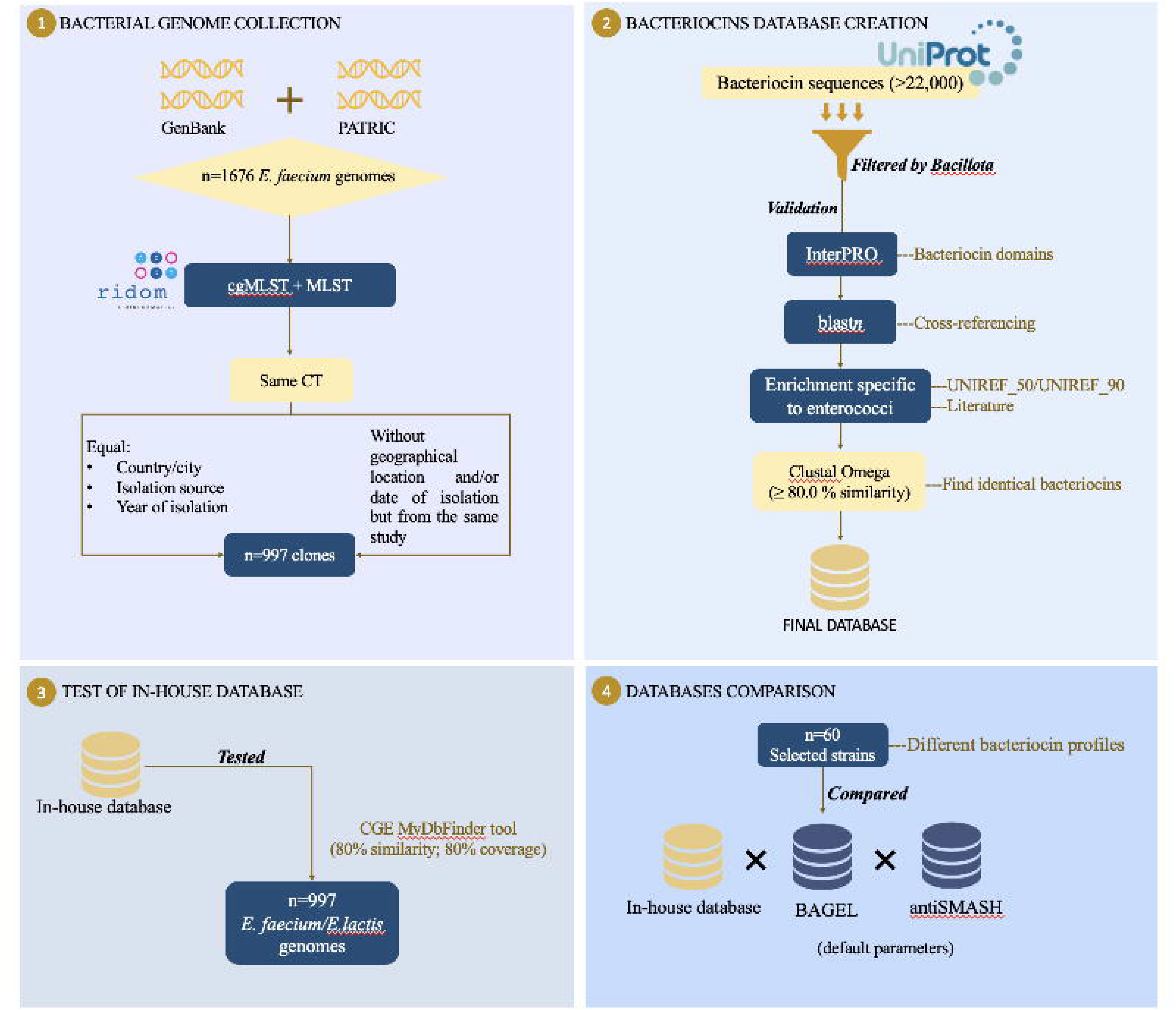
The database development and genomic analysis workflow. A) bacterial genome collection; B) bacteriocin database creation; C) bacteriocin analysis in enterococci genomes; and D) comparison of our in-house database with other available bacteriocin databases. Abbreviations: cgMLST - Core genome Multilocus Sequence Typing (cgMLST); MLST - Multilocus Sequence Typing; CT - cluster types; CGE – Center for Genomic Epidemiology.

The phylogenetic analysis of the 997 genomes was conducted using PATO software [11], and the resulting phylogenetic tree was visualised using iTOL [12]. To determine clonal relationships and distinguish isolates of *E. lactis,* we performed single nucleotide polymorphism (SNP) analysis, using PATO [11]. Furthermore, species confirmation for all 997 genomes was accomplished through *in silico* testing of the *gluP* gene sequences specific to *E. faecium* and *E. lactis* [13]. This involved employing a custom BLAST database and the makeblastdb tool, which is available in NCBI-BLAST v.2.14.0+.

### Creation of the bacteriocin database

Bacteriocin sequences (>22,000) were extracted from the Uniprot database and filtered by Bacillota. To validate these sequences as true bacteriocins, a multi-step approach was performed. This involved confirmation of bacteriocin domains using InterPro (for specific domains please see supplementary table S1), cross-referencing using blast*n* for both domains and sequence similarities, and relevant literature to incorporate known bacteriocins that may not have been initially identified in the database (e.g., *bac*AS48). Additionally, a targeted enrichment of bacteriocin sequences specific to enterococci was performed by leveraging the UNIREF_50/UNIREF_90 databases from Uniprot. The final set of bacteriocin sequences within the database was established through the application of Clustal Omega, where sequences exhibiting a similarity of ≥80.0% (as most genes are considered similar if they have a similarity ≥80.0%) were considered to belong to the same bacteriocin (Figure 1.B) [14]. Homologous bacteriocin sequences were eliminated from the database and bacteriocin sequences with identities between 80% and 99% were considered as the same bacteriocin but the sequences were maintained in the database in order to give more accurate identification. All bacteriocins were named using their UniRef100 code plus the name of the bacteriocin and in most cases (when this information was available) the bacterial strain they were isolated from.

### Whole-genome sequencing analysis of bacteriocins, antibiotic resistance genes, and plasmid replication initiation proteins

The detection of the bacteriocins present in the *E. faecium/E. lactis* genome*s* was performed by using our bacteriocin database in the CGE MyDbFinder tool (80% similarity and 80% coverage) (Figure 1.C). Antibiotic resistance genes and plasmid replication initiation proteins (*rep*) were screened using ResFinder 2.0 and PlasmidFinder 2.0 tools, respectively, available at the Centre for Genomic epidemiology (CGE; http://www.genomicepidemiology.org). Whenever available, genome metadata were utilized for antibiotic resistance annotation; otherwise, genotypes were evaluated using the ResFinder tool to identify antibiotic resistance genes and to categorize genomes into VRE (vancomycin-resistant enterococci), VSE (vancomycin-susceptible enterococci), AmpR (ampicillin-resistant enterococci) and AmpS (ampicillin-susceptible) groups.

### Databases comparison

The bacteriocin content identified in the enterococci isolates included in this study, using our in-house database, was cross-referenced with other specialized databases. Specifically, 60 selected strains representative of bacteriocin gene patterns diversity, according to our in-house database, were compared with BAGEL4 [8], a dedicated resource for bacteriocins, and antiSMASH [9], a tool designed for the analysis of secondary metabolite biosynthesis gene clusters in bacterial genomes. Both analyses were performed with the default parameters on the respective websites: BAGEL (http://bagel4.molgenrug.nl/) and antiSMASH (https://antismash.secondary metabolites.org/) (Figure 1.D).

### Statistical analysis

The Chi-squared test performed in Excel 2016 was used to determine associations (p≤0.05) between bacteriocin gene content and the two different enterococci species, sources, antibiotic resistance [mainly ampicillin (AmpR) and vancomycin (VanR) resistance] and plasmid replication initiation proteins.

## RESULTS

### Enterococcal species identification

The analysis based on the *gluP* sequence diversity of *E. faecium* and *E. lactis* revealed that the 997 *E. faecium* genomes available in the NCBI database indeed corresponded to 873 *E. faecium* and 124 *E. lactis.* While 114/124 belong to previously known *E. faecium* clade B, 10 of the 124 (≥98.71% identity with *gluP_E. lactis_* according to CGE MydBfinder analysis) belonged to *E. faecium* clade A1 and A2. When we checked the position of these 10 isolates within the SNP tree, they clearly belonged to *E. faecium* (Figure 2). These results indicate that in these strains *gluP* gene might be a hybrid between *E. faecium* and *E. lactis*. Considering the substantial genomic plasticity of *E. faecium* and its close relationship with *E. lactis*, we anticipate that this occurrence might be more frequent than initially anticipated [15]. Even though further analyses are needed in order to understand the *gluP* results obtained from these 10 strains, we classified them as *E. faecium* due to their positions in the SNP tree. Therefore, we analysed the genomes of 883 *E. faecium* and 114 *E. lactis*.

**Figure 2.**
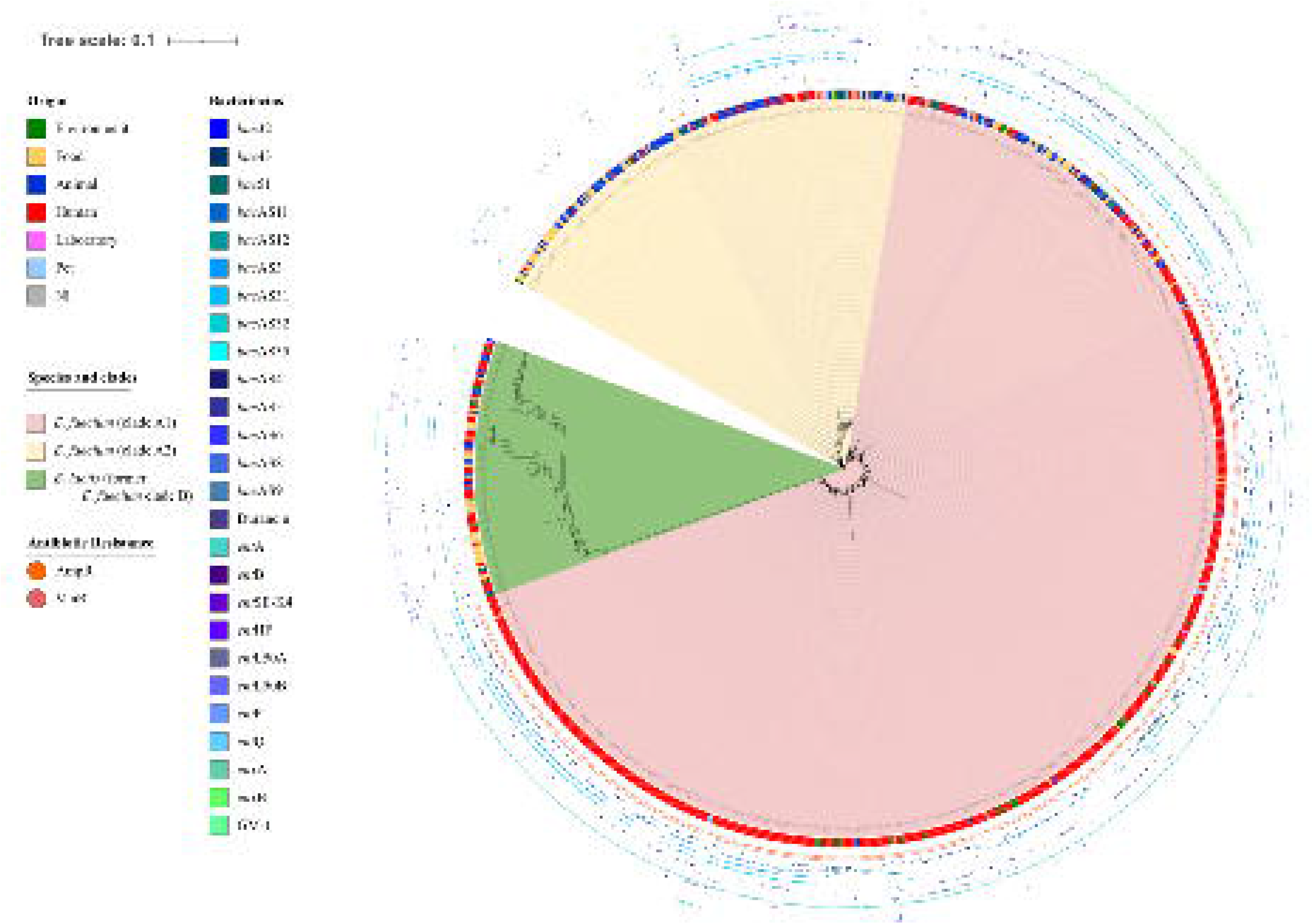
SNP tree of all *Enterococcus* genomes (n=997) analysed. The two isolates presenting genomes with a hybrid *glu*P gene are marked in red and bold.

### Overall characteristics of bacterial genomes

Enterococci genomes (n=997) were isolated from humans (n=612), animals (n=180), food products (n=130), environment (n=49), laboratory strains (n=7) and 19 had no information regarding source (Table S2). We have information on the year of isolation for 795 *E. faecium* and 89 *E. lactis*, which were collected between the years 1946 and 2019. The year with the highest number of represented strains was 2014 and the notable variations between different years align closely with extensive studies conducted in particular regions (Figure S1).

MLST analysis revealed the presence of 322 STs with 44 STs being new. Of the 322 STs 256 (27 new STs) were from *E. faecium* and 66 were from *E. lactis* (17 new STs). The most represented STs (Table S3) from *E. faecium* were those classically associated with nosocomial outbreaks such as ST17 (n=84), ST18 (n=60), ST203 (n=58), ST80 (n=53), ST412 (n=47), ST78 (n=31), ST117 (n=25), ST736 (n=18) and ST192 (n=17). There were other STs (Table S3) that have been previously associated with healthy humans, such as ST22 (n=17), and animals, as ST9 (n=15), ST32 (n=13), ST6 (n=13), ST54 (n=12), ST26 (n=10) and ST27 (n=10). For *E. lactis,* the most prevalent STs were those previously associated with food products, such as ST76 (n=11) and healthy humans, as ST94 (n=10) [16, 17]. cgMLST analysis revealed the presence of 847 CTs with the 740 CT belonging to *E. faecium* and 107 belonging to *E. lactis*. The most represented CT being CT128 (n=12), CT16 (n=11), CT24 (n=11) and CT722 (n=10) all belonging to *E. faecium*. Most CTs were represented by just one isolate (n=707) indicating a highly polyclonal bacterial collection (Table S3). The most represented countries in our genome collection were United States of America (USA, n=259), Australia (n=84), New Zealand (n=68), The Netherlands (n=54), United Kingdom (UK, n=50), Denmark (n=42), Canada (n=41), France (n=28), Russia (n=25), Sweden (n=25), Germany (n=14), Brazil (n=13) and South Korea (n=10) (Table S4). The country of origin was not reported for 45 isolates.

### Bacteriocin database and its application to *E. faecium* and *E. lactis* genomes

An in-house database of 76 Bacillota’s bacteriocins - 217 bacteriocin sequences; 40 new bacteriocins (<80% homology with other known bacteriocins as well as presenting known bacteriocin domains) with named “*bac*ASX” except for *bac*AS48, which has been previously described - belonging to classes I, II (subclasses IIa-IId) and III, was created (Table S1, fasta file bacteriocins). The most represented genera were *Enterococcus* (n=169/217), *Bacillus* (n=14/217), *Lactobacillus* (n=6/217) and *Lactococcus* (n=6/217) among other less represented genera from the Bacillota phylum (e.g. *Carnobacterium*, *Paenibacillus*, *Garciella* among others, Table S1). Such overrepresentation of the *Enterococcus* genus can be attributed to the higher number of genomes deposited at the NCBI and UNIPROT at the time of database creation (1,541 *versus* e.g., 336 *Lactobacillus* or 124 *Lactococcus*) as well as the high number of hypothetical proteins these genomes include, the manual enrichment (sequences added manually by the curator) in enterococcal sequences, and the naturally great production of bacteriocins by enterococci [2]. For these reasons, our database is only suitable for analysis of enterococci genomes and needs to be updated as more genomes are available.

Of the 76 bacteriocins listed in the database, 26 were found in the 997 *E. faecium* and *E. lactis* genomes included in this study. The results revealed that the predominant bacteriocins among *E. faecium* belong to subclass IIa and also subclass IIb (Table 1). The most prevalent bacteriocins were *ent*A (73.1%, 729/997, subclass IIa), followed by *bac*AS3 (38.7%, 386/997, subclass IIb), *bac*AS32 (38.5%, 384/997, subclass IId), *bac*AS11 (22.7%, 226/997, subclass IIc), *bac*43 (18.6%, 185/997, subclass IIa), *bac*AS5 (15.3%, 153/997, subclass IIb), *ent*B (subclass II-other) and *enx*A (12.1%, 121/997, subclass IIb), *bac*AS12 (10.9%, 109/997, subclass IIa), *enx*B (10.3%, 103/997, subclass IIb) and *ent*P (10.2%, 102/997, subclass IIa). The other 15 bacteriocins (5 subclass IIa, 1 subclass IIb, 4 subclass IId and 5 subclass II-other) were found in less than 10% of the genomes analysed (Figure 3 and Table S5). Differences in bacteriocin content were observed between *E. faecium* and *E. lactis* species (Figure 3). The temporal distribution of bacteriocins showed a consistent occurrence of different types during the whole period studied (Figure 4).

**Figure 3.**
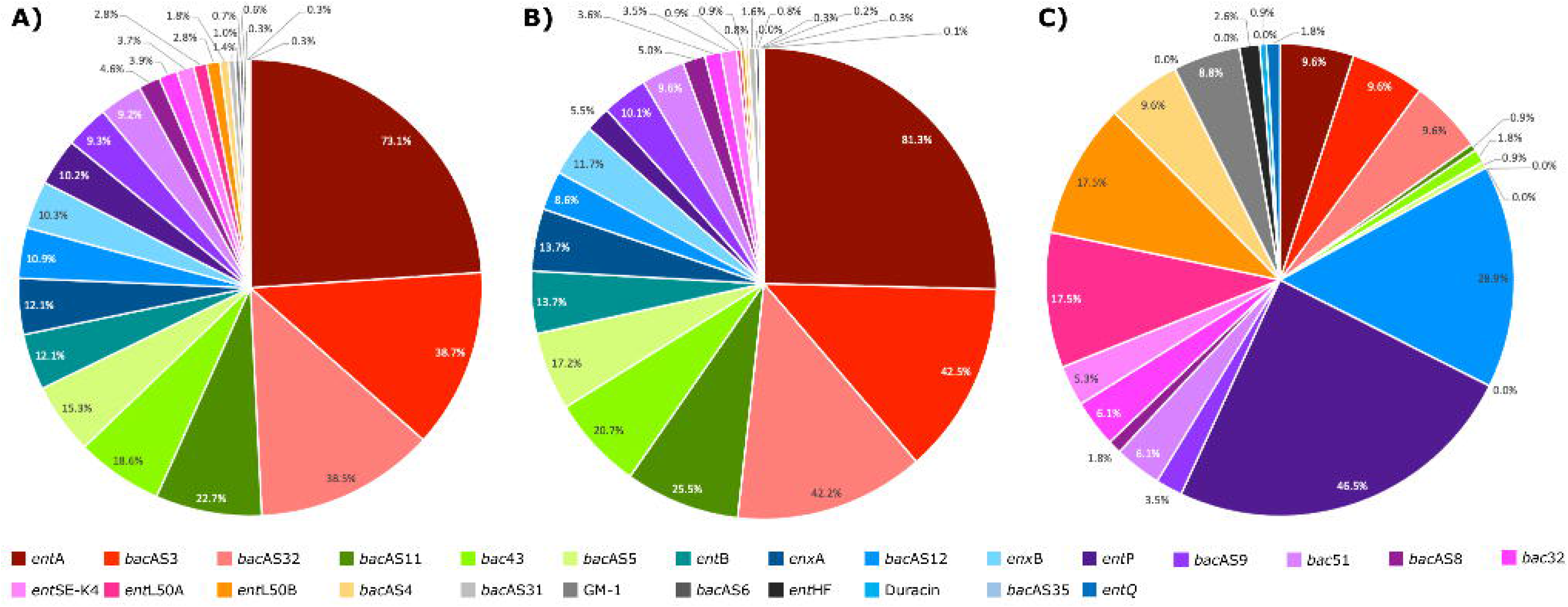
Comparative distribution of bacteriocins across A) all genomes, B) *E. faecium* genomes, and C) *E. lactis* genomes.

**Figure 4.**
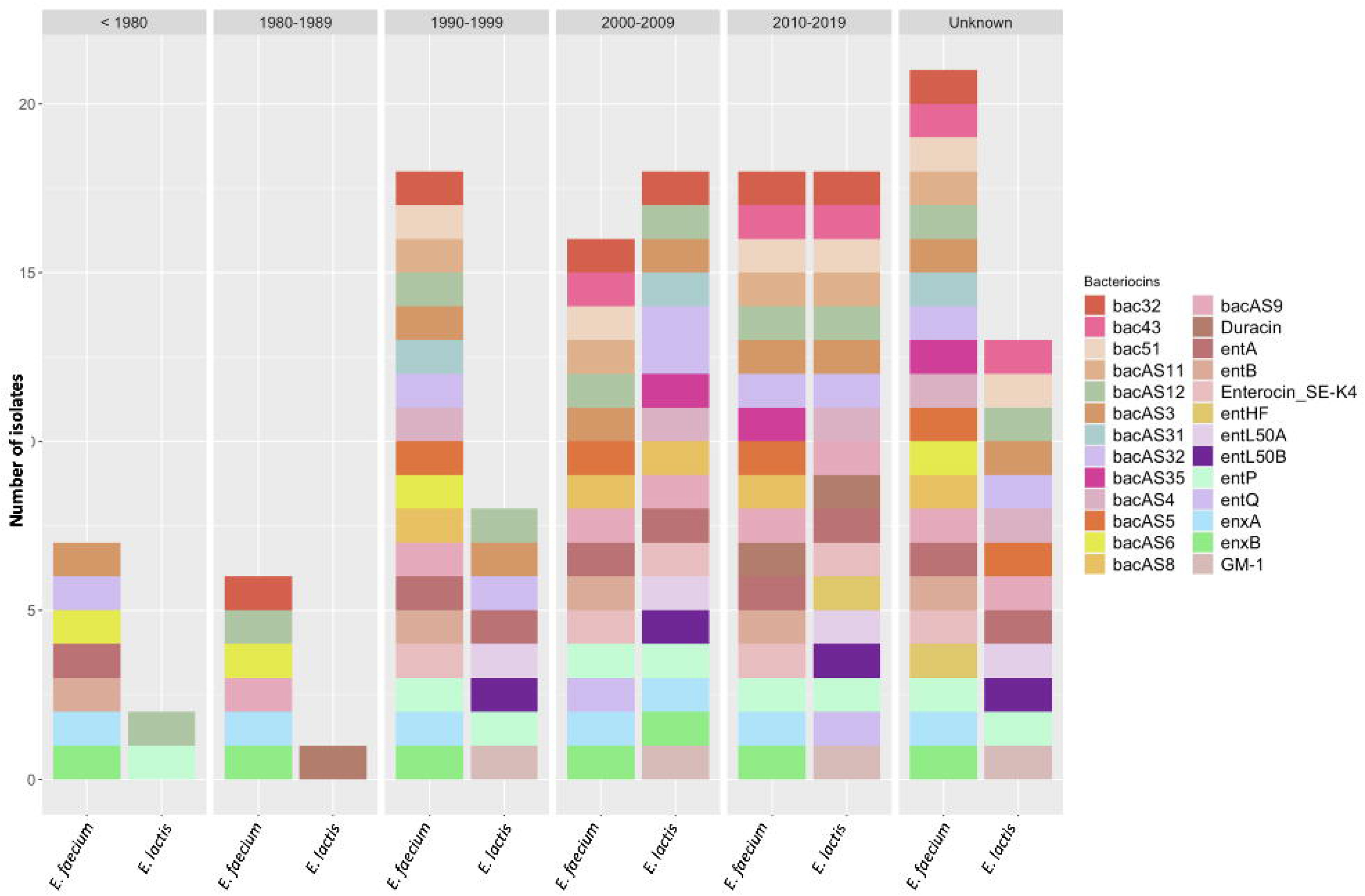
Comparative distribution of bacteriocins between *E. faecium* and *E. lactis* throughout the decades. Years before 1980 are grouped together due to low isolate numbers.

**Table 1.**
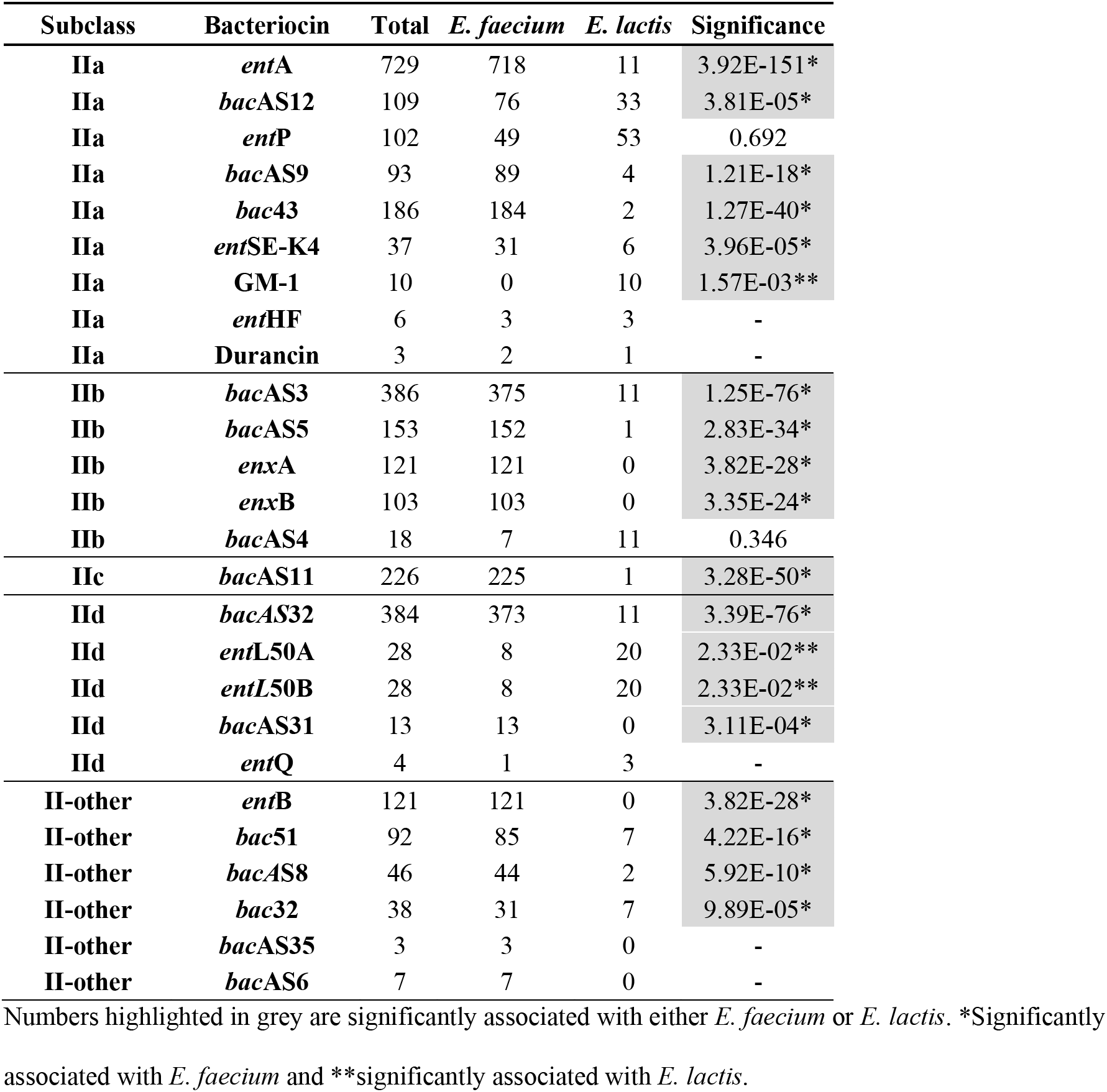
Number of bacteriocins present in all *Enterococcus*, *E. faecium* and *E. lactis* genomes and their associated with which *Enterococcus* species.

### Bacteriocin content in *E. faecium* and *E. lactis* genomes

The number of bacteriocins per genome spanned from 0-11 *bac* genes in the case of *E. faecium* and from 0-7 in the case of *E. lactis* species. Most *E. faecium* clinical isolates (406/433, 93.8%) carried 1-5 bacteriocins, while a significant portion of those sourced from animals (56/151, 37.1%) and foodstuffs (26/97, 26.8%) carried any bacteriocin. More than 7 bacteriocins were exceptionally observed among *E. faecium* genomes obtained from humans (n=4; two clinical, one healthy human, one unknown), animals (n=3; cow, rodent, horse) and food (n=1) sources (Figure 5).

**Figure 5.**
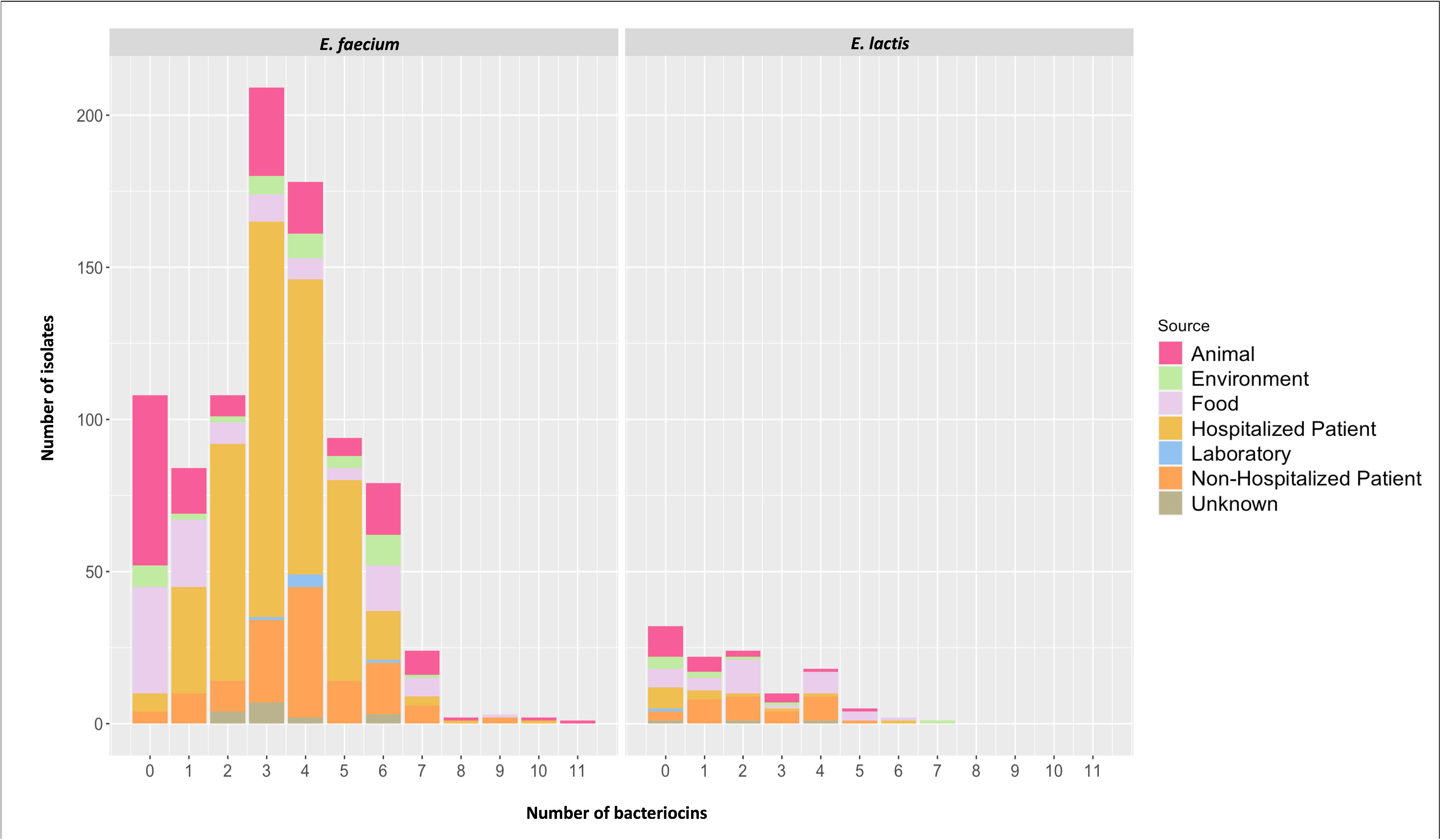
Number of isolates and their amount of bacteriocins (between 0 and 11 in *E. faecium* and between 0-7 in *E. lactis)* according to source and species.

The content of bacteriocin differs between *E. faecium* and *E. lactis* species (Figure 3). Out of the entire collection of bacteriocins, 16/26, corresponding to the ones present in more than 5% of isolates, were significantly associated with *E. faecium* (*ent*A, *ent*B, *enx*A, *enx*B, *bac*32, *bac*43, *bac*51, etc.; Figure 6; Table S5) while only three were positively associated with *E. lactis* (GM-1, *ent*L50A, *ent*L50B). Six bacteriocins (*ent*B, *enx*A, *enx*B, *bac*AS6, *bac*AS31 and *bac*AS35) were exclusively detected in *E. faecium* whereas only GM-1 was confined to *E. lactis,* predominantly from humans. Although this bacteriocin GM-1 was previously identified in *E. faecium* or *E. avium* strains [18], a recent analysis of bacteriocins in faecal samples of healthy humans only detected this GM-1 in *E. lactis* species [19]. According to the single available study evaluating the antimicrobial activity of GM-1, it exhibits a large spectrum against different Gram-positive (e.g., *Staphylococcus aureus, Bacillus* sp., *Listeria monocytogenes*) and Gram-negative (e.g., *Escherichia coli, Klebsiella pneumoniae, Pseudomonas aeruginosa*) species [3]. These observations suggest an adaption of GM-1 to the human host and implies a potential advantage for *E. lactis* as a prominent colonizer of the human gut. Some bacteriocins, as *ent*L50A/L50B, *bac*AS3/AS32 and *enx*A/B, were consistently found in pairs in both *E. faecium* and *E. lactis*.

**Figure 6.**
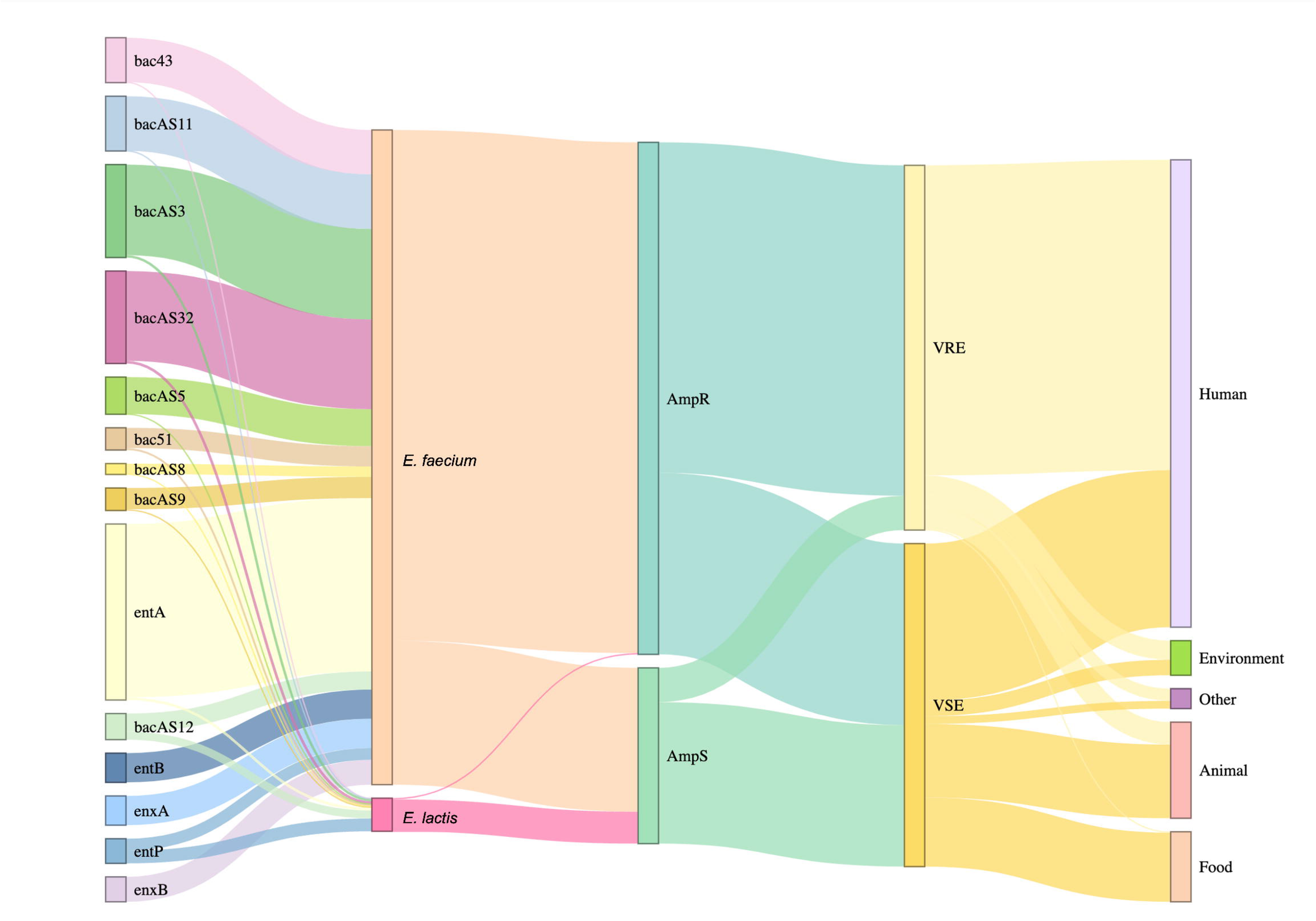
Sankey diagram illustrating bacteriocins’ distribution (present at least in 5% of the isolates) among *Enterococcus faecium* and *Enterococcus lactis* and associations with the presence/absence of antibiotic resistance genes as ampicillin (AmpR and AmpS) and vancomycin (VRE and VSE) and their respective sources. The diagram was constructed in R using the package networkD3 v0.4 [29]. Abbreviations: AmpR – Ampicillin resistant strains. AmpS – Ampicillin susceptible strains. VRE – Vancomycin resistant strains. VSE - Vancomycin susceptible strains.

### Host-specific adaptations revealed by source-based distribution of bacteriocins and association with antibiotic resistance genotypes

The epidemiological features of the strains containing bacteriocins is shown in Table S6. Some bacteriocins were enriched in *E. faecium* from humans (e.g., *ent*A, *ent*P, *ent*SE-K4, *bac*32, *bac*43, *bac*51, *bac*AS3/5/8/9/11/12/32), a statistically significant connection being noticed between *ent*A, *bac*43, *bacA*S5, and *bac*AS11 and strains from hospitalized patients (Figure 6; Table S6). Eight bacteriocins (*bac*AS4, *bac*AS12, *ent*B, *ent*P, *enx*A, *enx*B, *ent*L50A, *ent*L50B) were significantly associated with non-hospital sources, meaning that is expectedly very hard to find those *bac* genes in hospitalized patients ([3]; this study; Figure 6). The limited frequency of occurrence for *E. lactis* in most sources prevents us from deducing significant correlations these with sources.

We assessed the distribution of the different bacteriocins among genomes presenting different antibiotic resistance phenotypes (VRE *versus* VSE and AmpR *versus* AmpS) (Table S7; Figure 6). Six bacteriocins (*ent*A, *bac*43, *bac*51, *bac*AS5, *bac*AS9, *bac*AS11) could be significantly correlated with VRE, with four out of the six (*ent*A, *bac*43, *bac*AS5, and *bac*AS11) overlapping with those specifically associated with hospitalized patients and AmpR (Figure 6). *bac*AS9 and *bac*51 were both significantly associated with VRE and AmpR genomes, but not with hospitalized patients, and *bac*AS3 and *bac*AS32 (clustered together) were only linked to AmpR. Remarkably, among the 487 VRE genomes, only four belonged to *E. lactis*, and these four genomes exclusively contained the *ent*A bacteriocin among the six that exhibited a strong association with VRE. Among the aforementioned bacteriocins, those with available functional studies include *ent*A and *bac*43, mentioned in the previous section, and *bac*51, previously observed in clinical VREfm cases from Japan [20], has been further identified in this study and now expanded to clinical VREfm from various regions (Australia, Europe, USA, Brazil, Tunisia). Bac51 exhibits activity at least against different enterococcal species [3]. EntP, *ent*L50A, *ent*L50B, *bac*AS4 and *bac*AS31 were significantly associated with both AmpS and VSE, while *ent*B, *enx*A and *enx*B only with VSE (Figure 6). Interestingly, bacteriocins typically found in clinical isolates were strongly correlated with antibiotic resistance genes that are highly prevalent in the hospital environment, such as *ent*A and /or *bac*AS9 [*erm*(B)], *bac*43 [*erm*(B), *aph*(3’)-III, *aac*(6’)-Ie-*aph*(2’’)-Ia], *bac*AS5 [*erm*(B), *aph*(3’)-III], and *bac*AS11 [*erm*(B), *aph*(3’)-III, *van*A, *aac*(6’)-*aph*(2’’)] (Table S8).

### Plasmid distribution of bacteriocins

We also assessed if the bacteriocin distribution significantly correlated with the presence or absence of plasmid replication initiation proteins (*rep*). Several bacteriocins were associated with the presence of *rep*US15_pNB2354p1 (*ent*A, *bac*AS3, *bac*AS32, *bac*AS11, *bac*43, *bac*AS5, *bac*AS12, *ent*P, *bac*51 among others; Table S9) and *rep*2_pRE25 (*bac*43, *bac*AS11 and *bac*AS9) from the RepA_N and Inc18 plasmid families, respectively. The presence of *rep*US15_pNB2354p1 and *rep*2_pRE25 in a great number of isolates (670/997 and 440/997) could, up to a point, explain the association of these *rep* genes with so many bacteriocins since we did not assess the genome location of bacteriocins. Contrarily, several bacteriocins were associated with the absence of certain *rep* genes. For instance, *ent*A, *ent*B, *bac*AS3, *bac*AS32, *ent*P, *enx*A and *enx*B among others were associated with the absence of *rep*2_pRE25, *rep*US43_DOp1 and *rep*17_pRUM (Table S9). Several other *rep* genes were identified among the 997 enterococci isolates but their prevalence was not enough to allow a statistical analysis.

### Comparison of databases

We selected 60 isolates from our genome database, encompassing a wide variety bacteriocins, for comparative analysis between our in-house database and the antiSMASH and BAGEL databases. The antiSMASH database only positively detected the presence of the bacteriocin *ent*A in the 60 selected isolates, with a match in 29 isolates (Table 2). In contrast, our database identified *ent*A in 39 isolates. Notably, both databases identified *ent*A in 27 isolates, but 12 isolates exclusively tested positive for *ent*A according to our database, while 2 isolates were only positive for *ent*A in the antiSMASH database (Table 2).

**Table 2.**
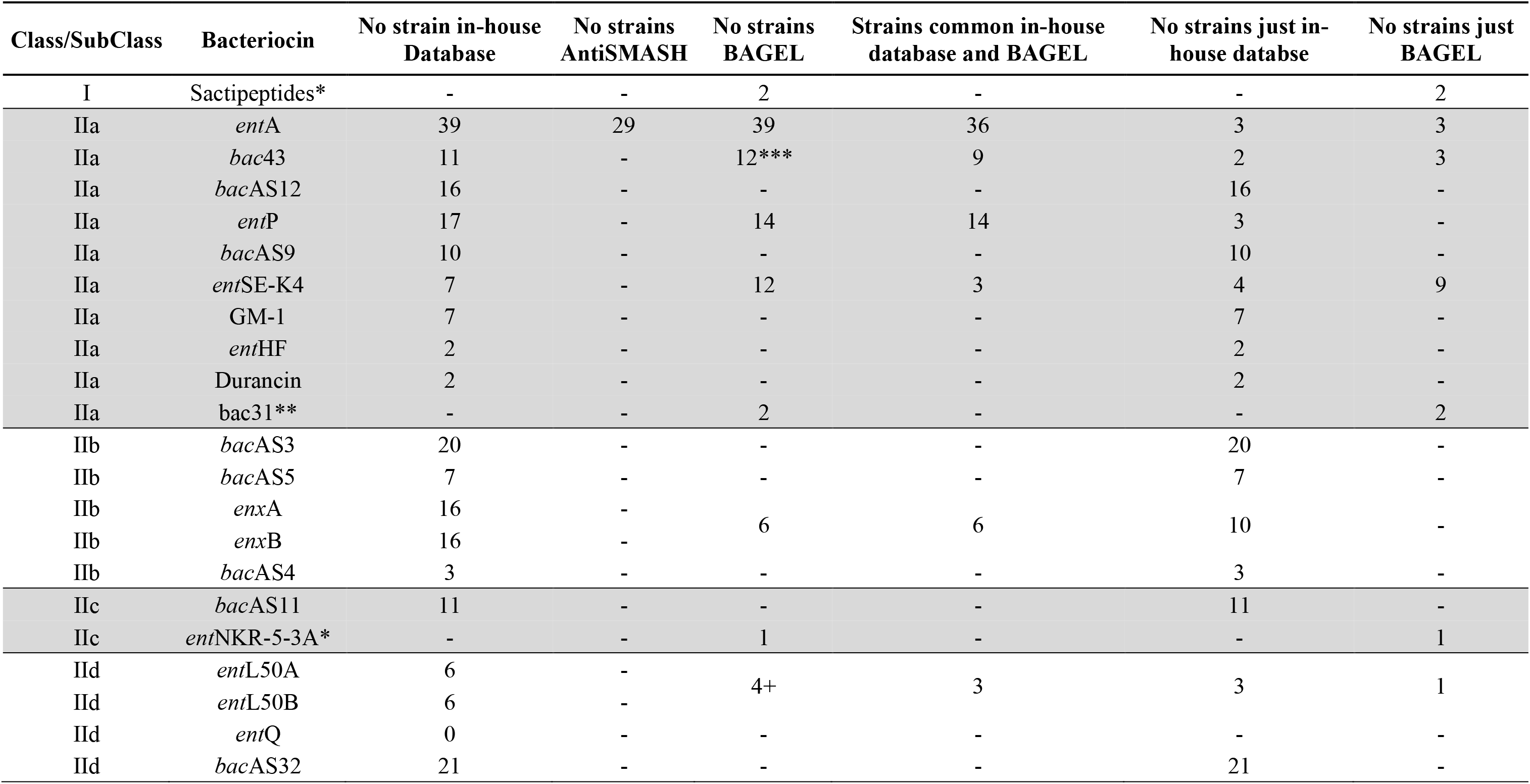
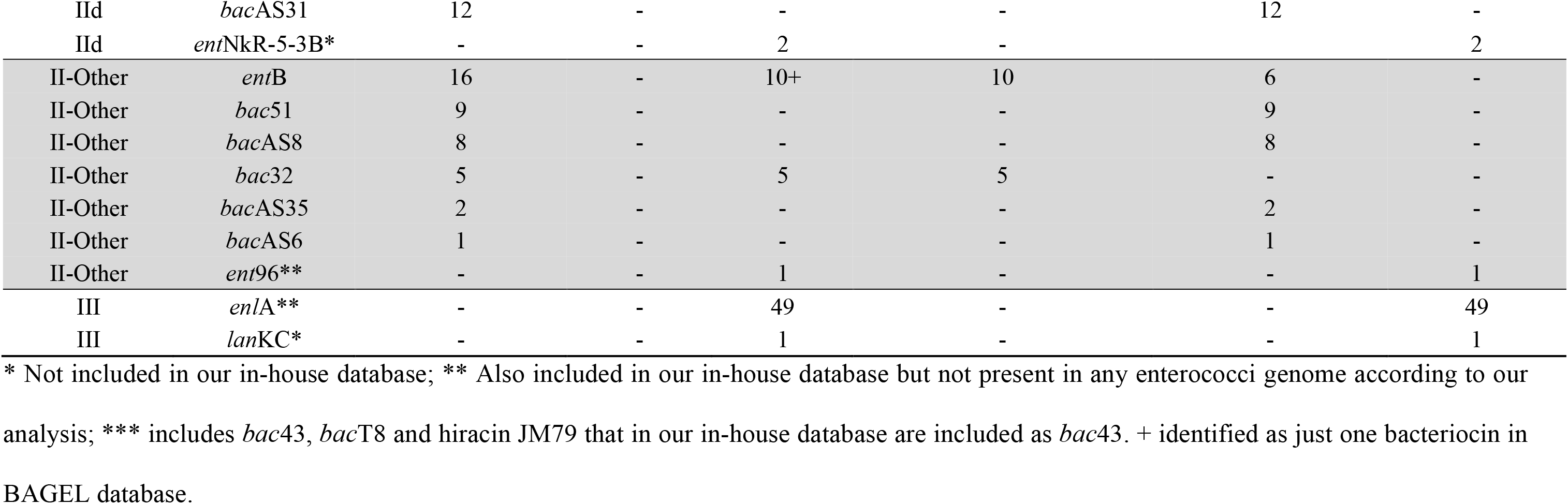
Bacteriocin results comparison between our in-house database and BAGEL and antiSMASH databases in 60 selected enterococci genomes.

The BAGEL database identified 18 bacteriocins in the 60 selected isolates while ours identified 26 bacteriocins (Table 2). Eleven of our database bacteriocins could not be detected in BAGEL and 4 of BAGEL bacteriocins were not detected by our database. Ten bacteriocins were detected by both BAGEL and our database: *ent*A, *bac*43, *ent*P, entSE-K4, *ent*B, *enx*A, *enx*B (BAGEL detects *enx*A and *enx*B as one unique bacteriocin), *bac*32, *ent*L50A and *ent*L50B (BAGEL detects *ent*L50A and *ent*L50B as one unique bacteriocin). In Table 2, it is evident that our proprietary database outperformed BAGEL in the detection of common bacteriocins across most isolates, with the notable exception of *ent*SE-K4, which BAGEL detected more frequently. The table provides a breakdown of the strains detected by both databases, exclusively by our database, or exclusively by BAGEL. Our database employs a strict cut-off of 80% identity and 80% coverage, as implemented in CGE. In contrast, BAGEL may exhibit lower identities in some cases (data not shown). This variance could explain instances where BAGEL detects bacteriocins not found by our database, contributing to discrepancies between the two resources. Notably, our database outperformed both BAGEL and antiSMASH, detecting a higher number of bacteriocins with greater accuracy.

## DISCUSSION

In this study, we analysed bacteriocin distribution patterns in *E. faecium* and *E. lactis* genomes using a homemade bacteriocin database in order to elucidate the contribution of bacteriocins to their adaptation in different environments/hosts and possible links to antibiotic resistance.

*E. faecium* genomes contained a diverse bacteriocin repertoire with evidence of host adaptation based on significant associations between bacteriocins such as *ent*A, *bac*43, *bac*AS5 and *bac*AS11, and human clinical isolates. Both *ent*A and *bac*43 belong to the class IIa of pediocin-like bacteriocins. *bac*43 has shown, in fact, a significant correlation with clinical *E. faecium,* including vancomycin-resistant *E. faecium* (VREfm), across different countries [17, 21], and exclusively sourced from human samples. The antimicrobial activity spectrum of *bac*43 has been scarcely explored but it is active at least against different enterococcal species and *L. monocytogenes* [3]. Although *ent*A has been observed in isolates from various sources, its strong association with human and clinical samples possibly indicates that, along with other bacteriocins, it may contribute to the ability of *E. faecium* to survive and compete in the gut. Enterocin A has been found to have inhibitory activity against a wide range of Gram-positive bacteria including staphylococci, listeriae and most lactic acid bacteria [3]. Enterocin B was significantly linked to animals and foodstuffs which corroborates previous studies showing its low occurrence in human clinical samples [3, 17].

Overall, several bacteriocins could be significantly associated with specific sources, potentially indicating host-specific adaptations. From an ecological perspective, bacteriocins may facilitate a producer’s introduction into an occupied niche as colonizing peptides, can directly inhibit competing strains or pathogens as antimicrobials or killing peptides or can serve as signalling peptides [22]. Although our approach did not take into account whether bacteriocin genes were integrated as complete operons or not, making it challenging to infer their expression, our study reinforces that *E. faecium* can harbour different bacteriocins per cell. Such arsenal can only provide a competitive advantage if its fitness benefit exceeds the metabolic cost of production, if it preserves essential mutualistic partner strains, and if it prevents major competitors from developing resistance [4]. Given that bacteriocin-producing *E. faecium* strains have the potential to beneficially modulate the indigenous microbiota [23] and in light of the observed variability in bacteriocin gene content within the species, as well as the specific adaptations seen in different sources and clades in this study, purified or partially purified bacteriocins offer substantial potential for targeting high-risk pathogenic clones effectively.

The association found between some bacteriocins and RepA_N plasmids is of note as this type of plasmid is prevalent among *E. faecium* from diverse sources and regions [17, 24–26] and has been also described in vancomycin-resistant (VanN) *E. lactis* [27]. Classically, Inc18 family plasmids and particularly *rep*2_pRE25 have been associated with VREfm as *van*A transposon is frequently located on these plasmids [17]. The bacteriocins associated with this type of *rep* gene have also been found to be associated with VRE genomes. Consequently, the presence of bacteriocins in these genomes may have been either co-selected with VanA plasmids or could, in conjunction with other factors, play a role in the stabilization of VRE within specific ecological niches [28]. On the other hand, some bacteriocins as *ent*A and *ent*B are associated with the absence of rep genes in the genome which might indicate their chromosomal location [3, 17] as described in previous studies. Bacteriocins have been often described as located on small mobilizable plasmids that seem to be usually cryptic (without known function) [3]. Therefore, it is crucial to give extra attention to these cryptic plasmids, especially since bacteriocin production can significantly contribute to plasmid maintenance, resembling the plasmid toxin-antitoxin systems and potentially facilitating their stable transmission across generations [28].

In summary, *E. faecium* clinical isolates were enriched in bacteriocins significantly linked to antibiotic resistance genotypes, including vancomycin resistance and Inc18 plasmids with *rep*2_pRE25. Together, these findings elucidate meaningful connections between bacteriocin determinants, antibiotic resistance, mobile genetic elements, and ecological origins in *E. faecium*. Such bacteriocin arsenal likely enhances the adaptability and competitive fitness of *E. faecium* in the nosocomial environment. Our analysis platform combining a novel tailored database, whole-genome sequencing, and epidemiological data provides a framework for elucidating bacteriocin landscapes in other organisms. Characterizing species-and strain-level differences in bacteriocin profiles may reveal determinants of ecological adaptation. Translating these discoveries could further inform strategies to exploit bacteriocins against high-risk clones.

## Supplementary material

**Table S1.** Description of our in-house bacteriocin database.

**Table S2.** Enterococci genomes distribution according to source and species.

**Table S3.** Enterococci genomes distribution according to species, source and cgMLST.

**Table S4.** Enterococci genomes distribution according to country of isolation and species.

**Table S5.** Bacteriocin frequency according to bacterial species.

**Table S6.** Bacteriocin frequency according to source.

**Table S7.** Bacteriocin frequency according to VanR/VanS and AmpR/AmpS.

**Table S8.** Bacteriocin frequency according to antibiotic resistance genotype.

**Table S9.** Bacteriocin frequency according to plasmid replication initiation protein. **Bacteriocin_database.fas.** Fasta sequences of all bacteriocins included in our in-house database.

**Figure S1**. Distribution of genomes retrieved from NCBI (n=997) per year of both *E. faecium* (n=883; yellow) and *E. lactis* (n=114; pink).

