## Supplementary figures and images for "Bacteriocin Distribution Patterns in *Enterococcus faecium* and *Enterococcus lactis:* Bioinformatic Analysis Using a Tailored Genomics Framework"

### Figure S1_Genomes_year_species.png

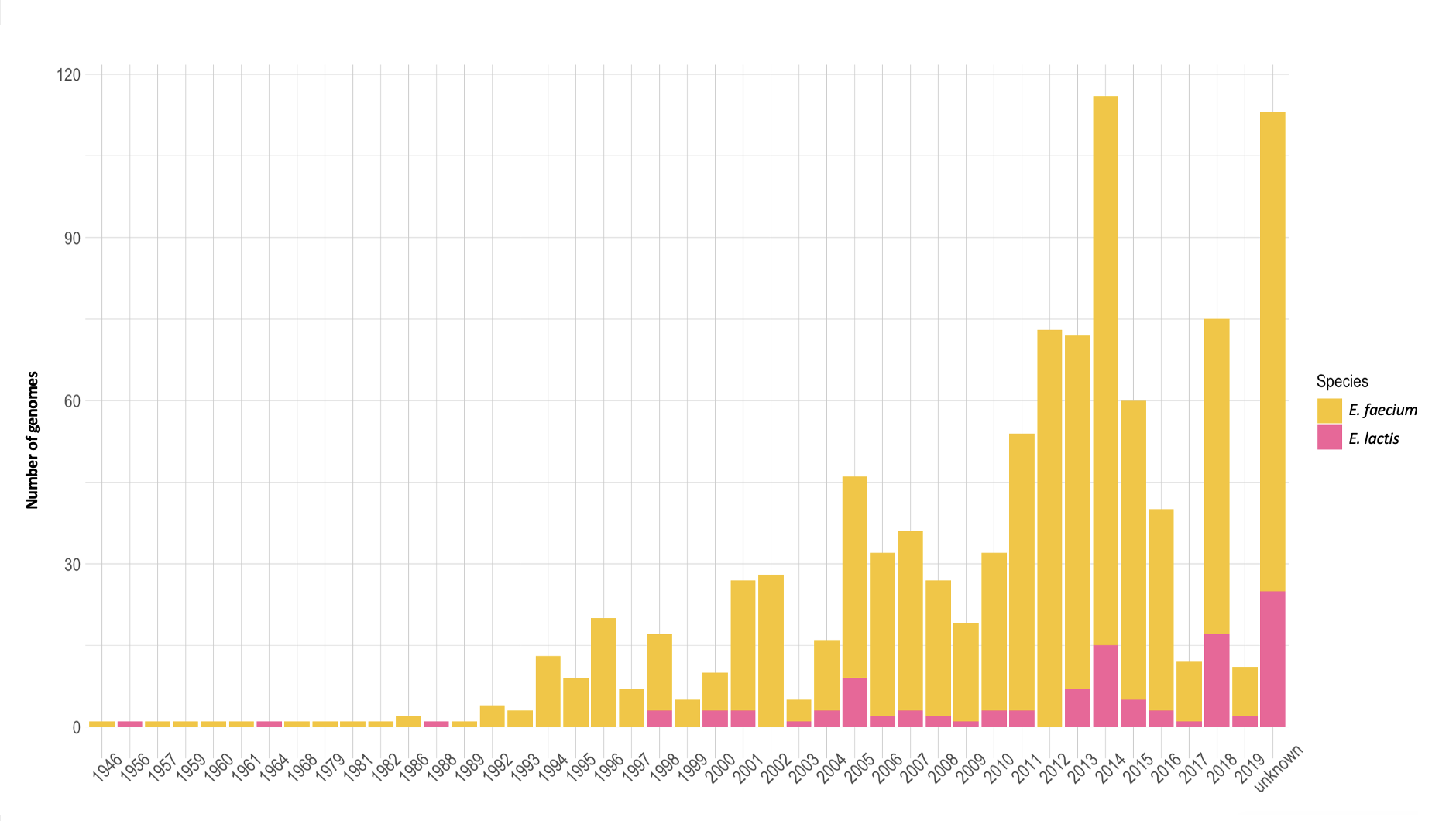
